# The BBSome assembly is spatially controlled by BBS1 and BBS4 in human cells

**DOI:** 10.1101/2020.03.20.000091

**Authors:** Avishek Prasai, Marketa Schmidt Cernohorska, Klara Ruppova, Veronika Niederlova, Monika Andelova, Peter Draber, Ondrej Stepanek, Martina Huranova

## Abstract

Bardet-Biedl Syndrome (BBS) is a pleiotropic ciliopathy caused by dysfunction of primary cilia. Most BBS patients carry mutations in one of eight genes encoding for subunits of a protein complex, BBSome, which mediates the trafficking of ciliary cargoes. Although, the structure of the BBSome has been resolved recently, the mechanism of assembly of this complicated complex in living cells is poorly understood. We generated a large library of human retinal epithelial cell lines deficient in particular BBSome subunit and expressing another subunit tagged with a fluorescent protein. We performed a comprehensive analysis of these cell lines using biochemical and microscopy approaches. Our data revealed that the BBSome formation is a sequential process including a step of the pre-BBSome assembly at pericentriolar satellites nucleated by BBS4, followed by the translocation of the BBSome into the ciliary base mediated by BBS1.

## Introduction

Bardet-Biedl Syndrome (BBS) is a multi-organ genetic disorder caused by the dysfunction of the primary cilia, microtubule-based sensory organelles. BBS is primarily characterized by retinopathy, polydactyly, genital and renal anomalies, obesity, and cognitive impairment [1]. Twenty-four causative BBS genes have been identified so far, eight of which form a stable protein complex called the BBSome [2, 3]. The BBSome is an adaptor protein complex consisting of BBS1, 2, 4, 5, 7, 8, 9, and 18, possessing structural similarities to coat/adaptor proteins involved in vesicular trafficking [3, 4]. The BBSome works in concert with the intraflagellar transport (IFT) machinery to facilitate the trafficking of particular transmembrane cargoes into and out of the primary cilia [5–9]. The role of the BBSome is pleiotropic as it mediates proper outcomes of multiple signaling pathways, including the Sonic Hedgehog signaling [10], leptin signaling [11], photoreceptor signaling [12], and neuronal signaling by G-protein-coupled receptors [13–15].

Mutations in any of the BBSome subunits can cause the BBS, suggesting that every subunit is essential for the complete BBSome function [2]. The structure of the BBSome was a longstanding enigma until recently, because the indirect approaches such as the yeast two-hybrid system [16], co-precipitation [17], co-expression of the individual subunits followed by low-resolution cryo-electron microscopy (EM) [18], and structural analysis of individual BBSome subunits [19, 20] provided only a very limited insight into the overall BBSome structure. Moreover, some authors proposed that BBS2, 7, and 9 form the BBSome core enabling the subsequent recruitment of other subunits [21], whereas others showed that BBS2 and BBS7 are dispensable for the assembly of the remaining six subunits [18, 22].

Three independent studies resolved the BBSome structure using cryo-EM with molecular modeling using the BBSome isolated from bovine retina [23] or using high-resolution cryo-EM analysis of bovine BBSome [24] or of human BBSome (lacking BBS2 and BBS7) expressed in insect cells [22]. These studies showed a high level of interconnectivity among the BBSome subunits, raising the question of how these BBSome proteins assemble in the cells.

Quantitative fluorescence microscopy techniques can reveal the formation and dynamics of functional protein complexes in living cells [25, 26]. In this study, we applied multiple microscopy and biochemical techniques to a large library of genetically engineered human RPE1 cells to describe spatially resolved steps of the BBSome formation in cells.

## Methods

### Antibodies

Mouse anti-acetylated tubulin was kindly provided by Dr. Vladimir Varga (IMG AS CR, Prague, CZ) and rabbit anti-BBS4 was kindly provided by Prof. Maxence Nachury (UCSF, CA, USA). Rabbit antibodies against BBS1 (ab222890) and acetylated tubulin (ab179484) were purchased from Abcam. Mouse antibodies against BBS2 (sc-365355), BBS5 (sc-515331), BBS8 (sc-271009), PCM-1 (sc-398365) and Ninein (sc-376420) were purchased from Santa Cruz Biotechnology. Rabbit antibody against BBS5 (14569-1-AP) was purchased from Proteintech. Rabbit antibodies against BBS7 (HPA044592), BBS9 (HPA021289), GFP (SAB4301138) and mouse antibody against actin (A1978) were purchased from Sigma-Aldrich. Rabbit antibody against actin (#4967) was purchased from Cell Signaling Technology. Rabbit antibody against NPHP1 (E-AB-61290) was purchased from Elabscience.

Secondary antibodies: anti-mouse Alexa Fluor 488 (Invitrogen, A11001, 1:1000), anti-rabbit Alexa Fluor 488 (Invitrogen, A11008, 1:1000), anti-mouse Alexa Fluor 555 (Invitrogen, A21422, 1:1000), anti-rabbit Alexa Fluor 555 (Invitrogen, A21428, 1:1000), anti-mouse Alexa Fluor 647 (Invitrogen, A21235, 1:1000) and anti-rabbit Alexa Fluor 647 (Invitrogen, A21245, 1:1000).

### Cell cultures

Immortalized human retinal pigment epithelium cells, hTERT-RPE-1 (ATCC, CRL-4000), henceforth RPE1, were kindly provided by Dr. Vladimir Varga (IMG AS CR, Prague, CZ) and Phoenix Ampho cells were kindly provided by Dr. Tomas Brdicka (IMG AS CR, Prague, CZ). Cells were cultured in complete Dulbecco’s Modified Eagle’s Media (DMEM) (Sigma, D6429-500mL) supplemented with 10% FBS (Gibco, 10270-106), 100 Units/ml penicillin (BB Pharma), 100 μg/ml streptomycin (Sigma Aldrich) and 40 μg/ml gentamicin (Sandoz) at 37°C and 5% CO_2_.

### Cloning, gene transfections and deletions

*BBS* ORFs were amplified from cDNA obtained from RPE1 cells, fused with the *SYFP2* coding region using recombinant PCR and cloned into pMSCV-Thy-IRES 1.1 vector (Clontech) using EcoRI/XhoI and ClaI restriction sites.

30 μg of plasmid DNA was transfected into Phoenix Ampho cells using the polyethylenimine to generate YFP-BBS retro-viruses. RPE1 cells were spinfected using 2 mL of virus aliquot in the presence of 2.4 μg/mL polybrene using standard protocols. RPE1 cells stably expressing YFP-tagged BBSome subunits were sorted using the 488 nm laser on FACSAria IIu (BD Biosciences).

*BBS* knockout cell lines were generated using the CRIPR/Cas9 approach. Single guided RNA (sgRNA) targeting *BBS* genes were designed using the web tool CHOPCHOP [27]. sgRNA was cloned into pSpCas9(BB)-2A-GFP (PX458) vector kindly provided by Feng Zhang (Addgene plasmid #48138) [28]. sgRNA sequence with PAM motif (3’ end) for respective *BBS* genes are listed below:

Human *BBS1*: GCAATGAGGCCAATTCGAAGTGG
Human *BBS2*: GTGGCCATAGGGCGCTACGACGG
Human *BBS4*: GGCTGAGGAGAGAGTCGCGACGG
Human *BBS7*: CAGTGTTCAAGACTTTACCCGGG
Human *BBS8*: GGCCGGTACCTGGTCATAAGGGG
Human *BBS9*: GATTGGTGGTCTACTATTCTGGG
Human *BBS18*: CAGAAGTGAAGTCAATGTTCCGG

RPE1 cells were transfected with PX458 vector with specific *BBS* sgRNA using polyethylenimine (Polysciences, Inc., 23966-2). After 48 h, cells expressing GFP were sorted as single cells in 96-well plates using the 488 nm laser on FACSAria IIu (BD Biosciences). Clones were then analyzed for the expression of target BBS proteins by immunoblotting and sequencing of DNA surrounding the sgRNA target site.

### Co-immunoprecipitation and western blotting

RPE1 cells were grown in 15 cm cell culture dishes and lysed in the lysis buffer (20mM HEPES at pH 7.5, 150mM NaCl, 2mM EDTA at pH 8, 0.5% Triton X-100) supplemented with protease inhibitor cocktail (complete, Roche, #05056489001). Lysates were cleared by centrifugation at 15,000 × g for 15 minutes at 4°C. Protein concentration was adjusted using the Pierce™ BCA Protein Assay Kit (Thermo Scientific). For co-immunoprecipitation assay, protein lysates were immunoprecipitated with anti-GFP antibody conjugated to protein A-coupled polyacrylamide beads (#53142 Thermo Scientific) for 2 h at 4°C. Beads were washed three times in the lysis buffer and co-precipitated proteins were denatured in 1× Laemmli buffer. For protein expression analysis, protein lysates were immediately denatured in reducing 4× Laemmli buffer. Denatured protein samples were analyzed by Western blotting following standard protocols. Membranes were probed with primary antibodies overnight at 4°C and secondary antibodies for 1 h at room temperature and developed using chemiluminescence immunoblot imaging system Azure c300 (Azure Biosystems, Inc.).

### Flow cytometry

RPE1 cells expressing the YFP-tagged subunits were cultivated in 24-well dishes, trypsinized and centrifuged at 1000 × g for 1 min. The pellets were washed in PBS and re-suspended in FACS buffer (2mM EDTA, 2% FBS and 0.1% sodium azide). Measurements were taken on Aurora™ (Cytek Biosciences). Geometric means of fluorescence intensity (GMFI) of YFP positive cells were obtained and used to examine the protein expression. Flow cytometry data was analyzed using FlowJo (BD).

### Immunofluorescence

RPE1 cells were cultured on 12 mm coverslips and serum starved for 24 h. Cells were fixed (4% formaldehyde) and permeabilized (0.2% Triton X-100) for 10 minutes. Blocking was done using 5% goat serum (Sigma, G6767-100ml) in PBS for 15 minutes and incubated with primary antibody (1% goat serum/PBS) and secondary antibody (PBS) for 1 h and 45 minutes, respectively in a wet chamber. The cells were washed after each step in PBS 3×. At last, the cells were washed in dH2O, air-dried and mounted using ProLong™ Gold antifade reagent with DAPI (Thermo Fisher Scientific).

### Fluorescence microscopy

Image acquisition for cilia length rescue assays was performed on the Delta Vision Core microscope using the oil immersion objective (Plan-Apochromat 60× NA 1.42) and filters for DAPI (435/48), FITC (523/36) and TRITC (576/89). Z-stacks were acquired at 1024 × 1024 pixel format and Z-steps of 0.2 microns. The cilia length was measured using the Fiji ImageJ.

Imaging of the YFP-tagged cell lines was performed using Zeiss Axioplan2 microscope equipped with an oil immersion objective (Plan-Apochromat Zeiss 100× NA 1.4) and filters for DAPI (435/48), YFP (559/38), TRITC (576/89) and Cy5 (632/22).

### Expansion microscopy and basal body length quantification

Protocol is based on [29]. RPE1 cells were cultured on 12 mm coverslips and serum starved for 24 h. Coverslips with cells were fixed with 4% formaldehyde/4% acrylamide in PBS overnight and then washed 2× with PBS. The gelation was performed by incubating coverslips face down with 45 μl of monomer solution (19% (wt/wt) sodium acrylate, 10% (wt/wt) acrylamide, 0.1% (wt/wt) N,N’-methylenbisacrylamide in PBS supplemented with 0.5% TEMED and 0.5% APS, prepared as described in [29] in a pre-cooled humid chamber. After 1 min on ice, chamber was incubated at 37°C in the dark for 30 min. Samples in the gel were denatured in denaturation buffer (200 mM SDS, 200 mM NaCl, 50 mM Tris in ddH2O) at 95°C for 4 h. Gels were expanded in ddH2O for 1 h until they reached full expansion factor (4.2×) and then cut into 1×1 cm pieces. Pieces of gel were incubated with primary antibodies diluted in 2% BSA in PBS overnight at RT. After staining, shrunk pieces of gel were incubated in ddH2O for 1 h, during which time they re-expanded. After reaching their original expanded factor, pieces of gel were incubated with secondary antibodies diluted in 2% BSA in PBS for 3 h at RT. Last expansion in ddH2O with exchange every 20 min was for 1 h until pieces of gel reached full size. Samples were imaged in 35 mm glass bottom dishes (CellVis, USA) pre-coated with poly-L-lysine. During imaging, gels were covered with ddH2O to prevent shrinking.

Expanded cells were imaged by confocal microscopy on Leica TCS SP8 using a 63× 1.4 NA oil objective with closed pinhole to 0.4 AU. Cilia and detail of basal bodies were acquired in Z-stacks at 0.1 μm stack size with pixel size 30-50 nm according to the cilia length. Images were computationally deconvolved using Huygens Professional software.

For basal body length, only those perfectly or nearly perfectly oriented longitudinally to image focal plane were selected for measurements. The line scan and plot profile tools in Fiji ImageJ were used to determine the width at the proximal end of basal body marked with Ac-tub antibody. Length of basal body was measured as a distance between proximal end (Ac-tub signal) and distal end stained by NPHP1 antibody. At least four independent experiments were measured in each condition (WT, n=56; *BBS1* KO, n=56; in total). For quantification of basal body length, in every experiment, the average value of widths was compared to ideal basal body width in full expansion (4.2×) to define the difference of ideal to real expansion factor (note, the range of expansions was between 3.35× and 4.1×). Then, as the length is subject of variability, all lengths were standardized by recounting the difference to full 4.2 expansion factor in particular experiment, and second, the average length was counted.

### Fluorescence recovery after photobleaching, data processing, and analysis

Cells were seeded in glass-bottom 8-well chambers (CellVis, USA) and serum starved for 24-48 h to induce ciliogenesis. Prior to fluorescence recovery after photobleaching (FRAP) experiments, cells were washed and supplemented with HBSS media (+Ca and +Mg) with 20mM HEPES. FRAP measurements were performed using a Leica TCS SP8 confocal microscope equipped with an oil immersion objective HC Plan-Apochromat 63× NA 1.4 oil, CS2 at room temperature. Data acquisition was performed in 512×512 pixel format with pinhole 2.62 Airy, at a speed of 1000 Hz in bidirectional mode and 8-bits resolution. Photobleaching (0.3 s) was performed with a circular spot 1 μm in diameter (centrosome, pericentriolar satellites (PS)) or with a rectangle spot 1 μm in length (ciliary base and tip) at 100% intensity using a 20 mW 488 nm solid-state laser. Fluorescence recovery was monitored at low laser intensity (3-10%) at 0.26 s intervals until reaching the plateau of recovery, in total for 60 to 80 s after photobleaching. A total of 25–30 separate FRAP measurements were performed for each sample. All FRAP curves were normalized to fluorescence loss during acquisition following the subtraction of background fluorescence. Curve fitting was performed in GraphPad Prism software using the one-phase association fit. All individual curves were fitted at once to obtain the mean and 90% or 95% confidence intervals of the desired parameters, halftime T_½_ and the mobile fraction F_m_ (see Supplemental Tables 2 and 3). Since the initial recovery after bleaching at the centrosome/ basal body and PS was contaminated by the diffusion towards these compartments, for simplicity and to estimate only the halftime of the protein fraction bound at these compartments, we restricted fitting of our data to times t > 2 s as we did in our previous study [26].

### Fluorescence correlation spectroscopy, data processing, and analysis

Cells were seeded in glass-bottom 8-well chambers (CellVis, USA) and serum starved for 24 h to induce ciliogenesis. Prior to fluorescence correlation spectroscopy (FCS) experiments, cells were washed and supplemented with HBSS media (+Ca and +Mg) with 20mM HEPES. FCS measurements were performed using a Leica TCS SP8 SMD confocal microscope equipped with three SMD HyD detectors, FCS module HydraHarp400 (PicoQuant, Germany), water objective HC Plan-Apochromat 63× NA 1.2 and WLL 470-670 nm at 37°C. Samples were spot-illuminated with the 488 nm laser line with 10 μW laser power at the sample plane, using pinhole size 1 Airy unit and detector range 505-600 nm. Data acquisition was performed using simultaneously the LAS-X software for operating the microscope and Symphotime64 software (PicoQuant, Germany) to online visualize FCS curves and save TTTR data. Illumination spots were selected based on the reference images of the cells. Spots were selected in the cytoplasm depending on the cell line analyzed. Spots were positioned close to the PS in YFP-BBS4 expressing cells or *BBS1* KO cell lines. In the WT cells, spots were selected in the cytoplasm in close proximity to the primary cilia. At least, 10–30 FCS measurements with an acquisition time of 60 s were taken per each cell line.

The instrument was aligned and the detection volume was calibrated using 20 nM Abberior STAR 488 carboxylic acid (Abberior GmbH, Germany) calibration dye. TTTR offline data processing and fitting was performed in a custom made “TTTR Data Analysis” software. The analysis pipeline starts with splitting each recording into 10 equivalent parts, calculation of autocorrelation curve for each part, removal of outlier curves that contained signal from aggregates, estimation of standard deviation for each time point of the correlation curve and weighted non-linear least-square fitting by a standard one or two component anomalous 3D diffusion in a Gaussian shape detection volume model [30] with triplet state fraction [31]. The used equation corresponds to Equation 10 in Chapter 5 of the online FCS manual (Ahmed, S. *et al. Practical Manual For Fluorescence Microscopy Techniques. PicoQuant GmbH* (2016)). Initially, the correlation curves were all fitted with 1-component anomalous diffusion fit to obtain the average diffusion time τ (see Supplemental Table 1). The fits enabled us to normalize the amplitude of correlation curves to 1 for lag time 10 μs, which allowed us to visually examine the differences between the measurements obtained from WT and KO cell lines and thus evaluate the presence of any potential second component. Only in case of YFP-BBS4 in WT cells we observed such dynamic behavior and therefore for this data we performed fitting also with the 2-component anomalous diffusion model (see Fig. 2 C, inset).

**Figure 1.**
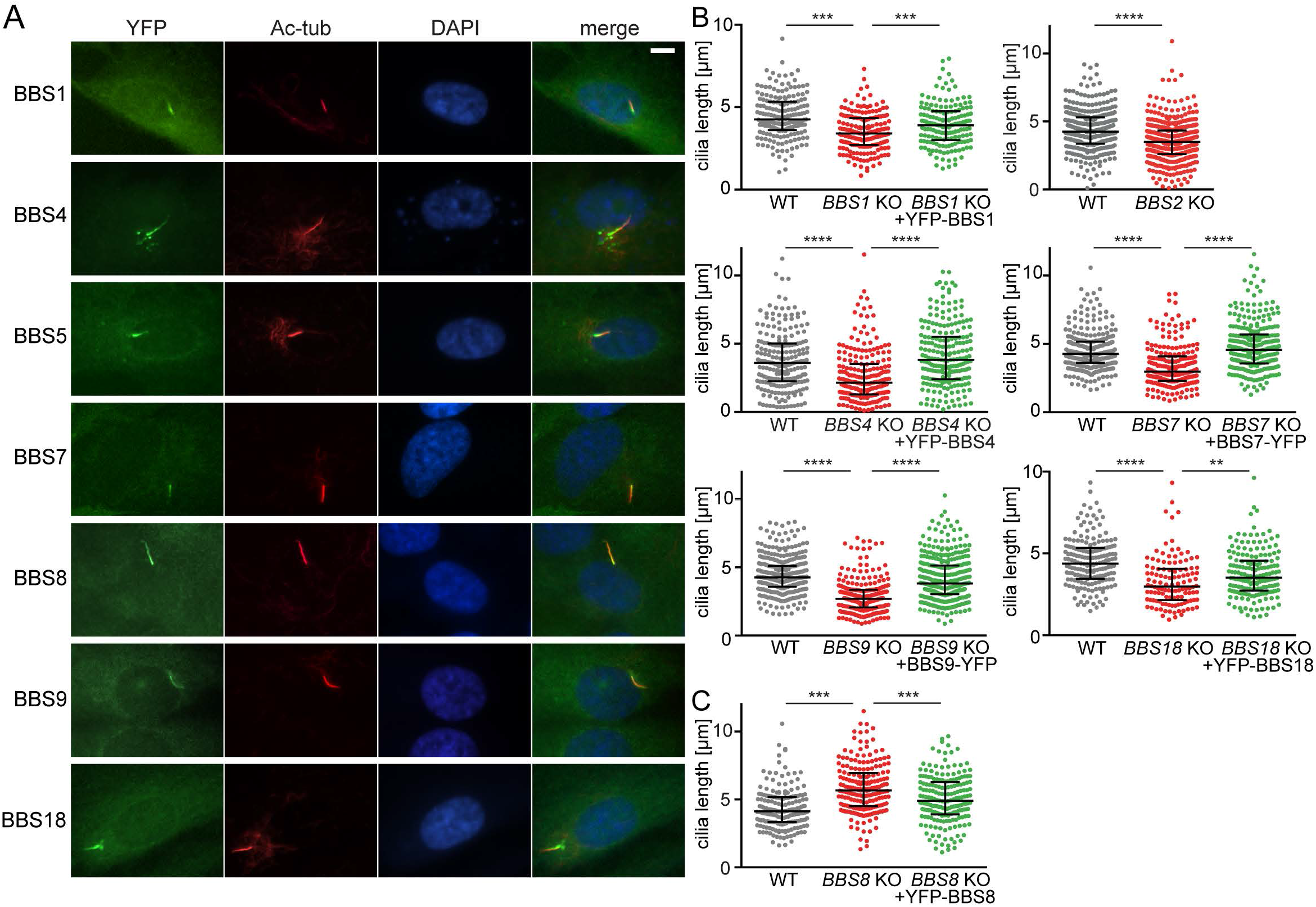
YFP-tagged BBSome subunits localize to the primary cilia and rescue the BBSome deficient cell lines. (A) YFP-tagged BBSome subunits localize to the ciliary base and primary cilia upon serum starvation in WT RPE1 cells. Antibody against acetylated tubulin (Ac-tub) was used to stain the primary cilia. Nucleus was stained with 4’,6-diamidino-2-phenylindole (DAPI). Scale bar, 5μm. (B) Plots show the comparison of ciliary length between WT, *BBS* KO and reconstituted *BBS* KO RPE1 cell lines. Ciliary length was rescued upon expression of the respective YFP-tagged BBSome subunit. Cilia length measurements were carried out using the Fiji ImageJ software. Medians with interquartile range from three independent experiments of n>100 are shown. Statistical significance was calculated using two-tailed Mann-Whitney test. **p<0.01, ***p<0.001 and ****p<0.0001. (C) Plot shows the comparison of ciliary length between WT, *BBS8* KO and reconstituted *BBS8* KO RPE1 cell lines. Ciliary length was restored to WT length upon expression of YFP-BBS8 in *BBS8* KO cell line. Cilia length measurements were carried out using the Fiji ImageJ software. Medians with interquartile range from three independent experiments of n>100 are shown. Statistical significance was calculated using two-tailed Mann-Whitney test. *** p<0.001.

**Figure 2.**
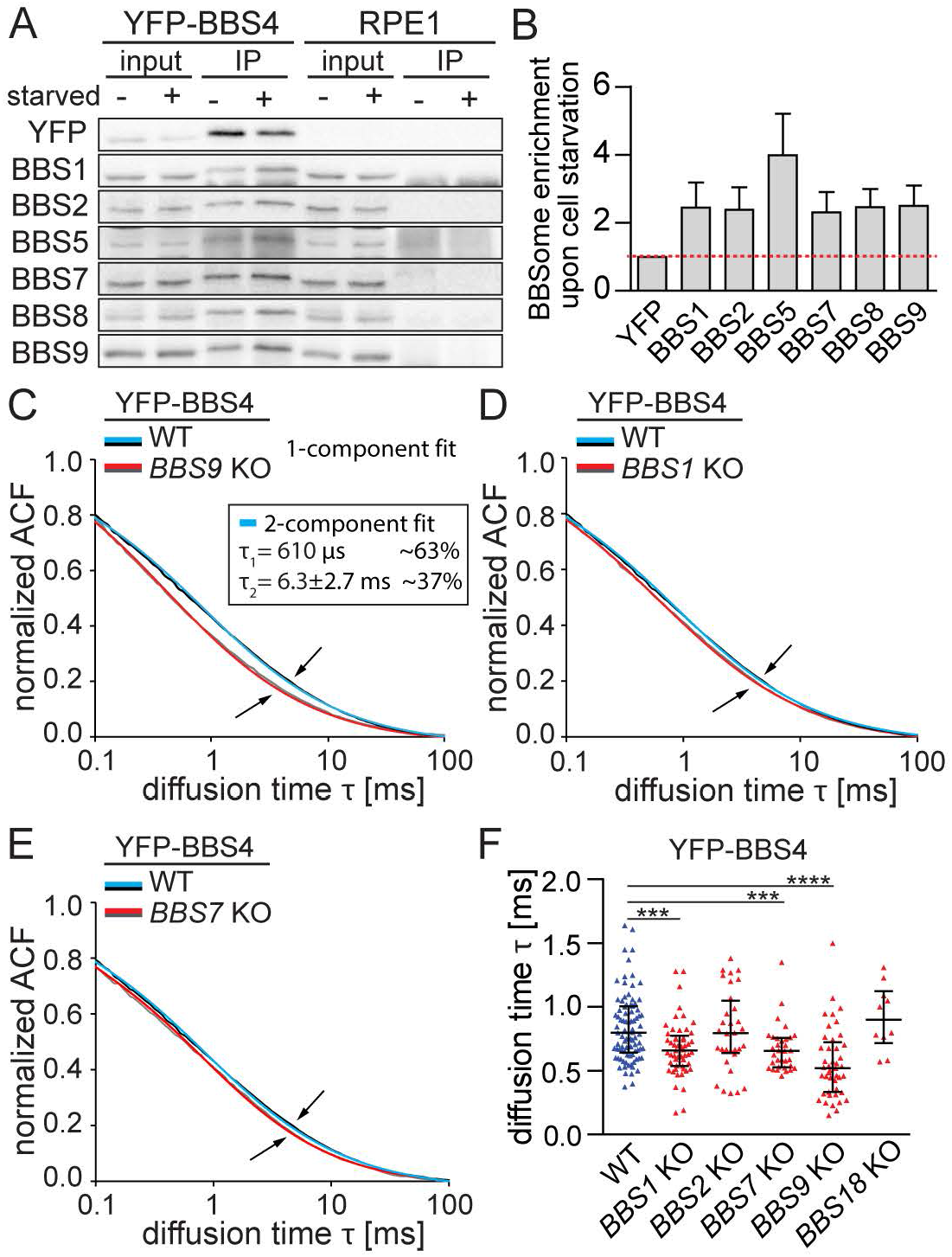
BBSome subcomplexes are present in the cytoplasm. (A) Co-immunoprecipitation of the BBSome subunits from YFP-BBS4 WT RPE1 cell lines using anti-GFP antibodies before and after 24 h serum starvation. Representative immunoblot out of three experiments is shown. (B) Quantification of the co-precipitated BBSome subunits was performed using the Fiji ImageJ software. Amount of the BBSome subunits was normalized to the YFP-BBS4 amount detected on the respective membrane. Mean with SD of three experiments is shown. (C, D, E) FCS measures the *in vivo* mobility of the BBSome subcomplexes in the cytoplasm. Plots show the autocorrelation functions (ACF) obtained from FCS measurements in the cytoplasm of YFP-BBS4 in WT and in *BBS9* KO (C), *BBS1* KO (D) and *BBS7* KO (E). The ACFs were fitted with one-component anomalous 3D diffusion model (Supplemental Table 1). The means of n>20 measurements are shown. Note the elevated mobility of YFP-BBS4 in the *BBS9* KO compared to *BBS1* KO and *BBS7* KO cells (arrows). YFP-BBS4 WT ACFs were additionally fitted by a two-component fit (inset). Diffusion time τ_1_ of the fast component was fixed to the mean diffusion time τ obtained from *BBS9* KO cells yielding the diffusion time τ_2_ of YFP-BBS4 engaged in a putative BBS4-BBS9 subcomplex. (F) Plot shows the diffusion times, τ, obtained by fitting the ACFs acquired from FCS measurements in YFP-BBS4 WT and *BBS* KO cell lines. Medians with interquartile range of n>10 are shown. Statistical significance was calculated using two-tailed Mann-Whitney test. ***p<0.001, ****p<0.0001

### Statistical analysis

Statistical analysis was performed using calculations built-in GraphPad Prism Version 5.04. All statistical tests were two-tailed. The statistical tests are indicated in the respective Figure legends.

## Results

### Generation of a cell line library for studying BBSome formation

To study the BBSome formation in cells, we initially established stable RPE1 cell lines expressing BBSome subunits fused with super yellow fluorescent protein SYFP2 (YFP) [32]. BBS1, BBS4, BBS8, and BBS18 were tagged at the N-termini, whereas the BBS7 and BBS9 were tagged at C-termini, because the N-terminal tag interfered with their function (data not shown). All YFP-tagged BBSome subunits localized to the primary cilia of RPE1 cells and showed weak diffuse signals throughout the cytoplasm (Fig. 1A).

In the next step, we generated a library of RPE1 cell lines deficient in *BBS1, BBS2, BBS4, BBS7, TTC8/BBS8, BBS9,* or *BBIP1/BBS18* using the CRISPR-Cas9 technology (Fig. S1A-B). Deletion of any of the BBSome subunits prevented the ciliary localization of YFP-tagged BBS1, BBS4, BBS5, BBS7, BBS8, BBS9, and BBS18 (Fig. S2A). Accordingly, endogenous BBS9 localized to primary cilia both in WT and reconstituted KO cell lines, but not in the KO cells, where it was rather diffuse throughout the cytoplasm (Fig. S3A). These data suggested that only the intact BBSome, but not BBSome intermediates or subunits alone could enter cilia.

Despite substantial overexpression of the YFP-tagged subunits over their endogenous counterparts (Fig. S3B), the fluorescence signal was still rather weak and mostly comparable in the respective WT and KO cell lines (Fig. S2A, S3C).

Cells deficient in BBS1, BBS4, BBS5, BBS7, BBS9, or BBS18 formed significantly shorter cilia in comparison to WT cells (Fig. 1B), corresponding to a previous observation in BBS4 deficient cells [33]. This phenotype was rescued by expressing YFP-tagged variants of the missing genes (Fig. 1B), documenting both that the cilia shortening was indeed caused by the gene deficiencies and that the YFP-tag does not interfere with the function of the BBSome subunits. Intriguingly, deletion of BBS8 resulted in prolonged cilia, which was reverted by the expression of YFP-BBS8 (Fig. 1C).

### BBSome subunits interact in the cytoplasm

Deficiency of one BBSome subunit reduced the cellular level of other subunits (Fig. S1A-B), indicating that the BBSome and/or BBSome intermediates are more stable than free individual subunits. In particular, deficiency in any of the subunits forming the proposed core of the BBSome (i.e., BBS2, BBS7, or BBS9) [21] substantially reduced the cellular levels of other subunits, whereas the absence of BBS1, BBS4, BBS8, or BBS18 had less dramatic effects on the stability of other subunits. The interdependence of the individual BBSome subunits suggests that they are present predominantly in the form of the BBSome or BBSome intermediates in WT cells.

We addressed whether ciliogenesis (induced by serum starvation) is coupled with de novo formation of the BBSome or whether the BBSome is pre-formed in non-ciliated cells (non-starved). BBS2, 5, 7, 8, and 9 co-immunoprecipitated with YFP-BBS4 in non-starved cells, showing that the BBSome assembles in non-ciliated cells (Fig. 2A). The serum starvation increased the amount of co-precipitated BBSome subunits ~2-fold, indicating that the BBSome formation is augmented upon ciliogenesis in RPE1 cells (Fig. 2A-B).

The presence of the BBSome in non-ciliated cells suggests that the BBSome or BBSome intermediates are present in the cytoplasm. Using FCS, we estimated the diffusion speed of YFP-tagged subunits in WT and KO cell lines (Supplemental Table 1). Because the large proteins and complexes diffuse slower than small complexes, these measurements reveal the information about the relative size of the respective complexes. We observed that the diffusion speed of YFP-BBS4 was significantly faster in *BBS9* KO cells than in WT cells, providing evidence that a fraction of BBS4 resides in a BBS9-dependent complex in the cytoplasm (Fig. 2C, F). We performed a two-component fitting of the WT data and found that the diffusion speed of this complex is about 10 times slower than the free YFP-BBS4 (Fig. 2C). Significant, but less pronounced effects were observed in *BBS1* KO and *BBS7* KO (Fig. 2D-F). These results show that cytoplasmic BBS4 exists in a form of a complex with BBS9 and likely other BBSome subunits. YFP-tagged BBSome subunits, other than BBS4, showed significant differences in their mobility between WT and KO cells only in some rare cases (Fig. S4A-D). It is possible that their over-expression masked the effect of BBSome disruption by favoring the monomeric forms (Fig. S3B-C). In case of YFP-BBS1, we could observe a mild loss in the slower fraction in *BBS7* or *BBS9* KO cells (Fig. S4B). Overall, these data imply the existence of heterogenic complexes of BBSome subunits in the cytoplasm.

### The pre-BBSome assembles at the pericentriolar satellites

BBS4 interacts with PCM-1 [34], which leads to its enrichment at the pericentriolar satellites (PS) both in ciliated and non-ciliated cells (Fig. S5A). BBS9 localizes to PS as well, albeit to a much lesser extent than BBS4 (Fig. 3A-B) indicating that a fraction of BBS9 and potentially other BBSome subunits reside at the PS in starved WT cells. Surprisingly, in *BBS1* KO cells, both the BBS4 and BBS9 were highly enriched at the PS (Fig. 3A-B). In addition, we observed that all other BBSome subunits (i.e., YFP-tagged BBS5, BBS7, BBS8, BBS9, and BBS18 and endogenous BBS2 and BBS9) localize to PS in *BBS1* KO cells (Fig. 3C, S5B). The enrichment of the BBSome subunits at the PS was dependent on BBS4, because it was not observed in BBS4-deficient cells and *BBS1/BBS4* DKO cells (Fig. 3D). To analyze the localization of BBS9 with higher resolution, we employed the expansion microscopy [29]. In line with the previous observations, BBS9 localized to the cilium in WT cells, localized to PS in *BBS1* KO cells, and was absent from the ciliary/centrosomal proximity in *BBS4* KO cells (Fig. 3E).

**Figure 3.**
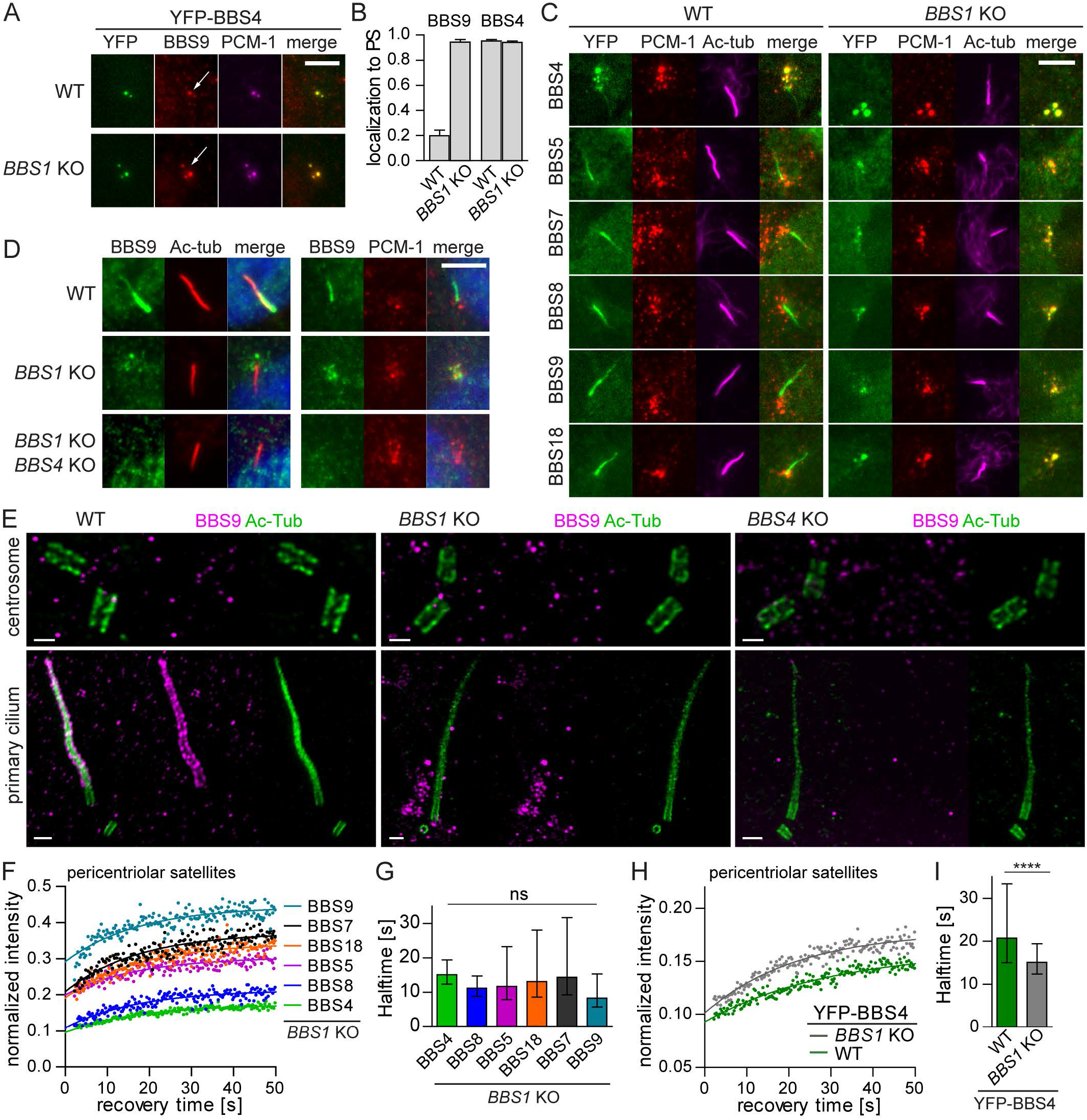
Loss of BBS1 stalls the pre-BBSome at the pericentriolar satellites. (A) YFP-BBS4 along with endogenous BBS9 localize to the PS both in WT and *BBS1* KO RPE1 cells (white arrows). BBS9 localization at the PS is pronounced in the *BBS1* KO RPE1 cells compared to WT RPE1 cells. Antibody against pericentriolar material-1 (PCM-1) was used to mark the PS. Scale bar, 5μm. (B) Quantification of localization of YFP-BBS4 and endogenous BBS9 to the PS in WT and *BBS1* KO RPE1 cells. Means and SD of two and three independent experiments respectively are shown. (C) YFP-tagged BBSome subunits localize to the primary cilia (Ac-tub) in WT RPE1 cells but are accumulated at the PS (PCM-1) in *BBS1* KO RPE1 cell lines. Scale bar, 5μm. (D) Endogenous BBS9 localizes to the primary cilia (Ac-tub) and the PS (PCM-1) in WT and *BBS1* KO RPE1 cells respectively. Deletion of BBS4 on top of BBS1 deficiency results in BBS9 dispersal form PS to the cytoplasm. Scale bar, 5μm. (E) Micrographs of BBS9 localization using expansion microscopy. BBS9 localizes to the cilia (Ac-tub) in WT RPE1 cells, around the ciliary base and at the PS in *BBS1* KO and is diffused in *BBS4* KO. Scale bar, 2μm. (F) FRAP analysis of the dynamic turnover of the YFP-tagged BBSome subunits at the PS in *BBS1* KO RPE1. Recovery curves were fitted at once using one phase association fit. Mean of 20-30 measurements from three independent experiments is shown. Estimated halftimes T1/2 and immobile fractions are shown in Supplemental Table 2. (G) Bar graph depicts recovery halftimes (s) of YFP-tagged BBSome subunits in *BBS1* KO cell lines obtained from FRAP analysis in F. Means of 20-30 measurements from three independent experiments are shown. Error bars represent the 95% confidence interval. p>0.05. (H) FRAP analysis of the dynamic turnover of the YFP-BBS4 at PS in WT and *BBS1* KO RPE1 cells. Recovery curves were fitted using one phase association fit. Mean of 20-30 measurements from three independent experiments is shown. Estimated halftimes T_½_ and immobile fractions are shown in Supplemental Table 2. (I) Bar graph depicts recovery halftimes (s) of YFP-BBS4 in WT and *BBS1* KO cell lines obtained from FRAP analysis in H. Means of 20-30 measurements from three independent experiments are shown. Error bars represent 95% confidence interval. The statistical difference was calculated by Mann-Whitney test. **** < 0.0001.

FRAP analysis revealed comparable recovery half-times for all YFP-tagged BBSome subunits in *BBS1* KO cells at the PS (Fig. 3F-G, Supplemental Table 2), possibly because they are predominantly engaged in a single complex. YFP-BBS4 showed the highest immobile fraction, which is consistent with its direct binding to the PS via PCM-1 (Fig. 3F). Most likely, BBS4 recruits other subunits to the PS leading to the formation of a pre-BBSome complex. The accumulation of the BBSome intermediates at the PS in *BBS1* KO cells results in the mobilization of BBS4 (Fig. 3H-I, Supplemental Table 2), indicating that the pre-BBSome binding to PS is less stable than binding of BBS4 alone. BBS1 is crucial for the completion of the full BBSome, which prevents the BBSome subunits, with the exception of BBS4, from the arrest at the PS.

### BBS1 facilitates BBSome translocation from the basal body to the cilium

As all BBSome subunits, BBS1 localizes to the cilia in WT cells (Fig. 1A). However, BBS1 localizes to the centrosome in non-ciliated cells and to the centrosome/basal body in some cells with short nascent cilia (Fig. 4A). Moreover, BBS1 localizes to the centrosome/basal body in *BBS4* KO cells, whereas other BBSome subunits show diffuse localization in the cytoplasm of these cells (Fig. 4B). Deficiency of BBS4 increased the turnover of BBS1 at the centrosome/basal body (Fig. 4C-D), suggesting that the interaction of BBS1 with the pre-BBSome stabilizes BBS1 at the centrosome/basal body, possibly directing the whole complex towards the cilium.

**Figure 4.**
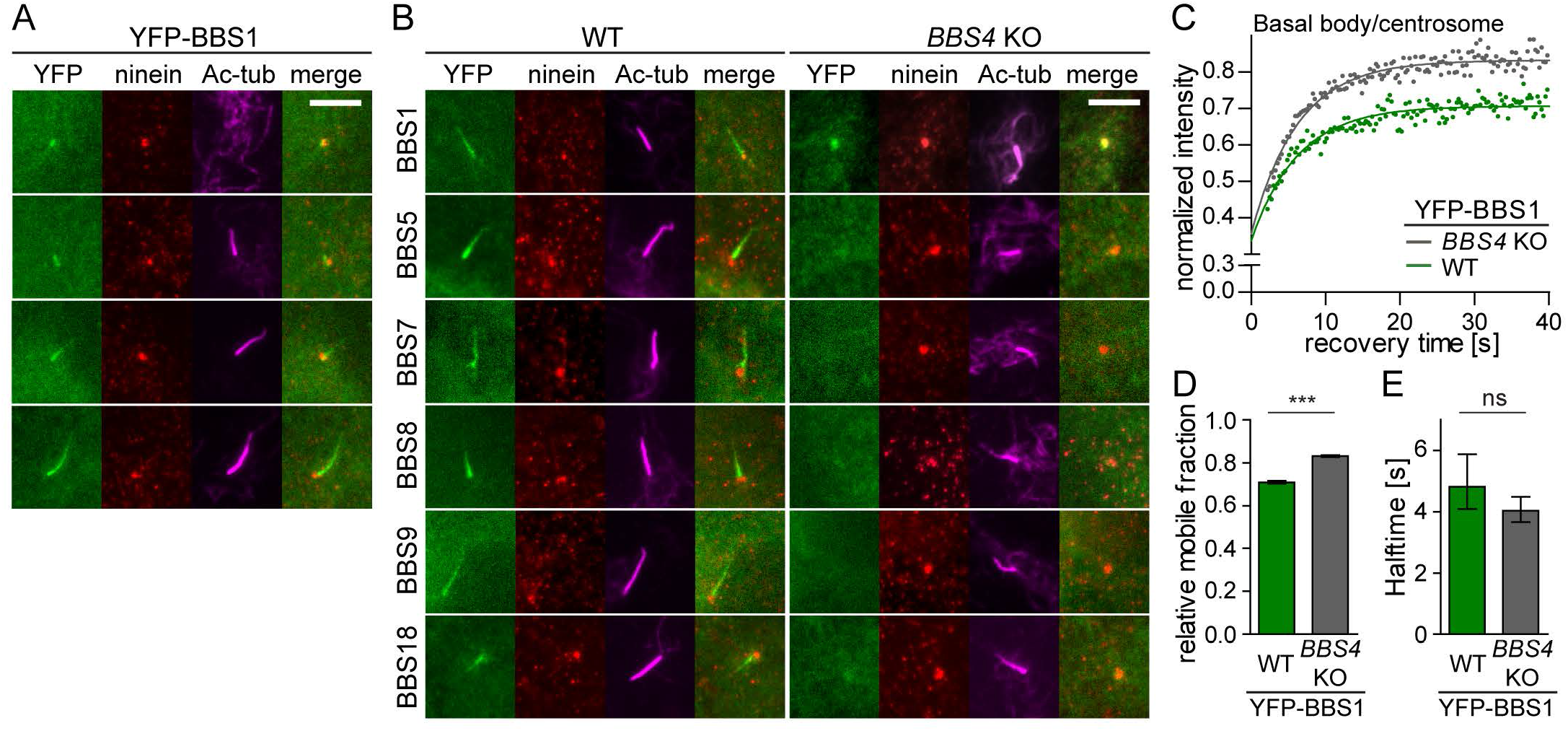
BBS1 acts as a gatekeeper at the basal body to facilitate BBSome completion and its entry to cilia. (A) YFP-tagged BBS1 transits through the centrosome into the ciliary base and the cilia in WT RPE1 cells. Ninein and acetylated tubulin staining visualizes the centrosome and cilia respectively. Scale bar, 5μm. (B) YFP-tagged BBSome subunits localize to the primary cilia (Ac-tub) in WT RPE1 cell lines. YFP-BBS1 localizes to the centrosome (Ninein) in the *BBS4* KO cells, whereas, the other YFP-tagged BBSome subunits are diffused in the cytoplasm. Scale bar, 5μm. (C) FRAP analysis of the dynamic turnover of the YFP-BBS1 at the centrosome or basal body in WT and *BBS4* KO RPE1 cells. Recovery curves were fitted using one phase association fit and the means of 20-30 FRAP measurements from three independent experiments are shown. Estimated halftimes T_½_ and immobile fractions are shown in Supplemental Table 2. (D) Bar graph depicts the relative mobile fraction of YFP-BBS1 in WT and *BBS4* KO cells extracted from FRAP analysis in C. YFP-BBS1 has a greater mobile fraction in *BBS4* KO than in WT RPE1 cells. Means of 20-30 measurements from three independent experiments are shown. Error bars represent the 95% confidence interval. ***p<0.001. (E) Bar graph depicts the recovery halftime of YFP-BBS1 in WT and *BBS4* KO cells obtained from FRAP analysis C. Means of 20-30 measurements from three independent experiments are shown. Error bars represent the 95% confidence interval. p>0.5.

We performed the analysis of the dynamic behavior of the BBSome subunits at the ciliary tip and the basal body and the transition zone using FRAP (Fig. 5A-C, Supplemental Table 3). If the subunits are incorporated in the whole BBSome quantitatively, their recovery rate should be comparable. However, BBS1 behaved differently from the other subunits, because it was very mobile at the base of the cilium (Fig. 5C, S5D). Indeed, BBS1 was the only subunit which was more dynamic in the ciliary base than in the ciliary tip (Fig. 5D, S5C-D). These data indicate that BBS1 exists in two forms at the ciliary base, as a monomeric protein with fast turnover between the ciliary base and the cytoplasm and as a part of the BBSome, whereas other BBSome subunits are recruited to the ciliary base only in the form of the complete BBSome complex.

**Figure 5.**
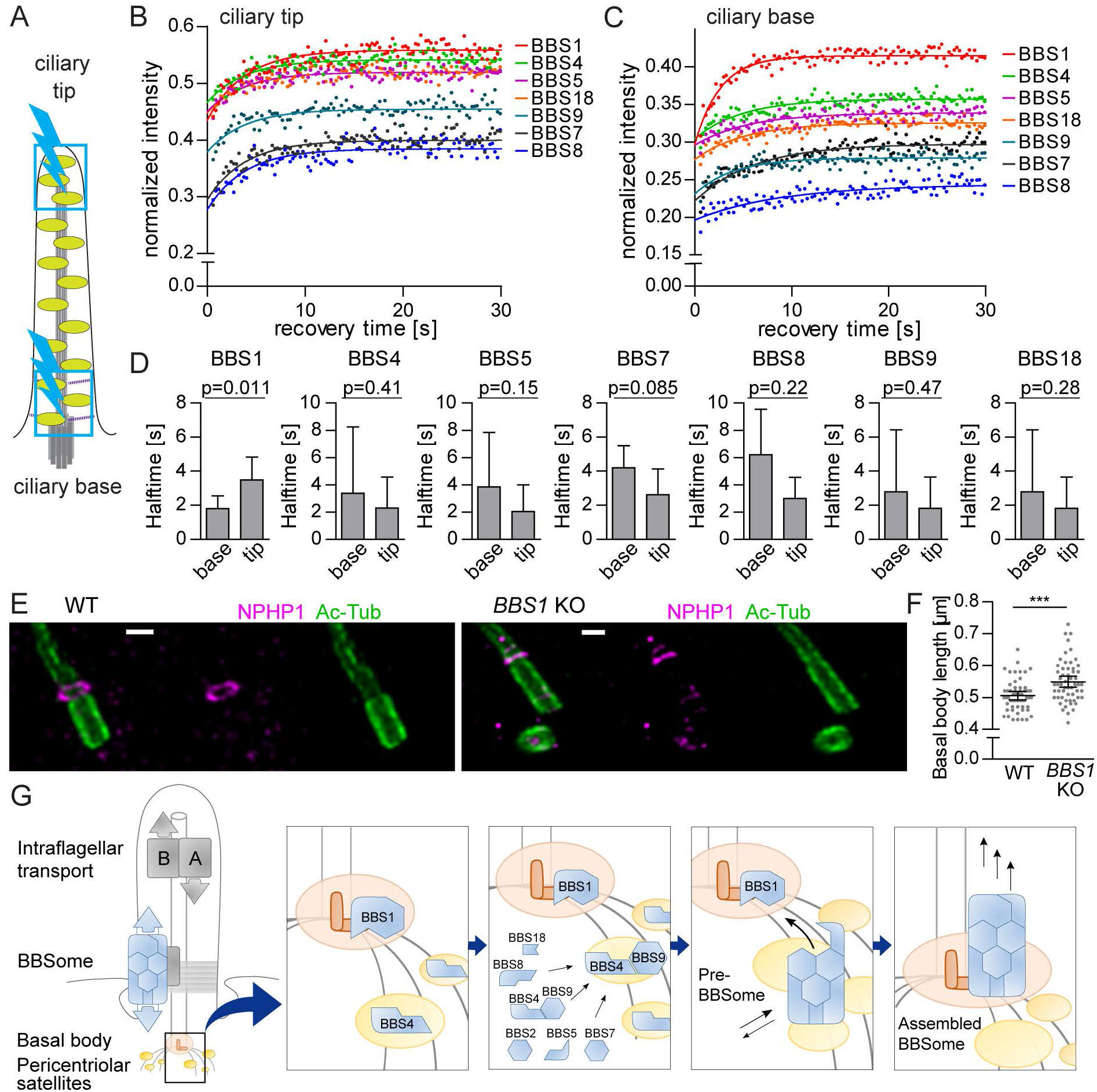
BBS1 has a distinct dynamic behavior at ciliary base. (A) Schematic representation of the FRAP experiment in the primary cilia. Ciliary base and tip (blue boxes) were bleached for 0.3 s using a 20 mW 488 nm solid-state laser at 100% intensity (blue lightning bolt). (B, C) FRAP analysis of the dynamic turnover of the YFP-tagged BBSome subunits at the ciliary base (B) and tip (C). FRAP curves were fitted with one phase association fit. Means of 20-30 measurements from three independent experiment are shown. Estimated halftimes T_½_ and immobile fractions are shown in Supplemental Table 3. (D) Bar graphs depict the recovery halftimes of BBSome subunits at the ciliary tip and base. Means of 20-30 measurements from three independent experiments are shown. Error bars represent the 90% confidence interval. (E) Micrographs of basal body (Ac-tub) and transition zone (NPHP1) staining in WT and *BBS1* KO RPE1 cells using expansion microscopy. Scale bar, 2μm. (F) Plot of the basal body lengths quantified based on the expansion microscopy performed in WT and *BBS1* KO cell lines in E. *BBS1* KO cells have significantly longer basal body compared to WT. Statistical significance was calculated using two-tailed Mann-Whitney test. Means with SD of n=56 are shown. ***p<0.001. (G) Schematic model of the spatially resolved formation of the BBSome in human cells. BBS4 is a resident protein at PS, while BBS1 resides at the basal body. BBS4 associates with BBS9 in the cytoplasm, which directs the BBSome subunits to the PS to assemble the pre-BBSome. Pre-BBSome is stabilized by BBS1 at the basal body resulting in the full BBSome competent to enter the cilia.

Interestingly, the absence of BBS1 not only stalls BBSome at the PS, but alters the structure of the ciliary base resulting in slightly prolonged basal body and transition zone (Fig. 5E-F), suggesting a role of the BBS1 in the organization of the ciliary base.

Based on our data, we propose a model of the BBSome formation in cells with the main roles for BBS4 and BBS1 in the spatial regulation of the full complex assembly (Fig. 5G). BBS4 resides on PS and via its association with BBS9, it recruits other BBSome subunits to form the pre-BBSome complex. BBS1 localizes to the basal body and guides the pre-BBSome to the ciliary base and subsequently to the cilium.

## Discussion

Because the BBSome consists of 8 subunits, it has been proposed that it assembles in a step-wise manner [21]. However, in vitro experiments provided contradictory data concerning the sequence of the individual steps [18, 21, 22]. Although the resolution of the structure of the BBSome complex represented a break-through in the field [22–24], these studies could not reveal how the BBSome assembles in living cells, including the spatial regulation of individual steps. We generated a library of 64 RPE1-derived cell lines using a combination of CRISPR/Cas9 knockouts of individual BBSome subunits and retroviral expression of YFP-tagged subunits. The only misses in the library were BBS5 KO and BBS2-YFP, because we were unsuccessful in their preparation. We used these lines to address the localization of individual BBSome subunits in WT cells and in cells lacking another subunit. The signal from YFP-tagged proteins was complemented with the detection of endogenous BBS9 and BBS2 using immunofluorescence. Moreover, we used two microscopy techniques, FCS and FRAP, to elucidate the dynamics of individual subunits in our cell lines. Altogether, our data enabled us to propose a model of the BBSome formation in living cells.

It has been shown that some BBSome subunits stabilize each other in cells [21, 35]. Our systematic analysis demonstrated that actually all BBSome subunits are interdependent. Deficiency in any of the subunits (with the exception of BBS5 which was not tested) leads to the substantial decrease of the cellular levels of other subunits, indicating that only a minor fraction of the BBSome subunit molecules are monomeric. Deficiency of BBS2, BBS7, or BBS9 had the most pronounced effects on the abundance of other subunits, suggesting that these three structurally related proteins form the core of the BBSome in vivo [21]. We showed that BBSome is readily assembled in non-ciliated cells and ciliogenesis only modestly augments the BBSome formation. Our FCS approach detected some direct or indirect cytoplasmic interactions of BBSome subunits, most notably BBS4-BBS9. Altogether our data show that BBSome and/or its intermediates are present in the cytoplasm and that the cilia is not required for the BBSome formation.

The PS organize proteins involved in the centrosome maintenance and ciliogenesis [36, 37]. It has been shown previously that BBS4 localizes to the PS, where it interacts with resident proteins PCM-1, Cep290, and AZI1 [38–40]. Our data indicated that BBS4 recruits other BBSome subunits to the PS, where a pre-BBSome complex is formed. Absence of PCM-1 disrupts the PS and reduces ciliogenesis [3], but does not prevent the ciliary localization of the BBS4 [40]. However, other PS proteins, Cep72 and Cep290, mediate ciliary localization of BBS4 and BBS8, suggesting that they orchestrate the pre-BBSome formation [40]. Our data show that the pre-BBSome is released from the PS and targeted to the basal body via BBS1, the only subunit that shows an intrinsic affinity for the centrosome/basal body.

Depletion of AZI1 leads to enhanced localization of BBS4 into the cilium and restores the ciliary localization of BBS9 in the absence of BBS5, partially in the absence of BBS2 and BBS8, but not in the absence of BBS1 [38]. Although these experiments used RNA interference which usually does not lead to the complete loss of the target protein, the data are in line with the fact that BBS1 is essential for the ciliary BBSome targeting. AZI1 might negatively regulate the release of the pre-BBSome from the PS. Interestingly, deficiency of AZI1 in zebrafish leads to the typical BBS symptoms [38], suggesting that the role of AZI1 is important for the BBSome function. LZTFL1 is a BBSome-interacting partner showing a similar loss-of-function phenotype to AZI1, including the ciliary localization of the incomplete BBSome [41] and the development of BBS symptoms, in this case in humans and mice [42, 43]. It is tempting to speculate that AZI1 and/or LZTFL1 mediate a quality control mechanism blocking the release of incomplete pre-BBSome from the PS. As BBS5 is a relatively peripheral subunit of the BBSome [22–24] that is sub-stoichiometrically incorporated into the complex in vitro [22], a control mechanism checking the incorporation of BBS5 into the pre-BBSome in cells seems to be plausible.

Earlier studies suggested that the sequestration of the BBS4 at the PS might deplete the pool of BBS4 accessible for the BBSome formation [38, 40]. However, the experimental data presented in these studies are consistent with our model of the pre-BBSome assembly at the PS and its subsequent release. Moreover, the putative quality control function of LZTFL1 and AZI1 might explain why the depletion of these proteins causes a seemingly paradoxical coincidence of enhanced ciliary localization of BBSome subunits as well as the induction of BBS-like phenotypes. Last, but not least, it would be difficult to reconcile the model of BBS4 sequestration at the PS with our observation that all BBSome subunits are strongly enriched at the PS in the absence of BBS1.

Our model proposes that BBS1 is incorporated into the BBSome as the last subunit, because of its affinity to the centrosome/basal body. However, we cannot exclude that the last step of the BBSome formation is not the incorporation of BBS1 itself, but another BBS1-dependent event. It could be a conformational change in the BBSome, caused by a BBS1-interacting GTPase, BBS3/ARL6 [9, 19, 24]. In any case, BBS3 seems to be involved in the translocation of the (pre-)BBSome from the PS to the ciliary base, as the BBSome-BBS3-GTP does not co-purify with PCM-1 [3, 9] and BBS3 was proposed to mediate the interaction of the BBSome with ciliary membranes [9, 24]. BBS1 has been shown to interact with RABIN8, a guanine-exchange factor for RAB8, which is involved in vesicular trafficking and might regulate BBSome targeting to the ciliary base [3]. Overall, it seems likely that BBS1 facilitates an energy-consuming conformational change of the complex that eventually confines the BBSome in the cilia.

Chaperonins BBS6, BBS10, and BBS12 assist the formation of the BBSome [21, 44] and the loss of any of them leads to the BBS [45–47]. Because the deficiency in BBS1 or BBS3 generally presents with a less severe BBS disease than the deficiency in BBS2, BBS7, or BBS9 core proteins or in the BBS chaperonins [2], it is probable that the chaperonin complex assists any of the initial steps of the BBSome formation as suggested previously [21] rather than the BBS1-dependent translocation into the ciliary base. However, this should be experimentally addressed in further studies.

Overall, our study reveals that the BBSome formation is a sequential process with spatially compartmentalized steps. The formation of the pre-BBSome is nucleated by BBS4 at the PS, whereas BBS1 drives the translocation of the BBSome to the ciliary base. It shall be addressed in follow-up studies whether some of the multiple reported causative mutations [2] interfere with the process of the BBSome assembly.

## Acknowledgement

We thank Dr. Vladimir Varga for sharing the reagents and the Zeiss microscope. We thank Dr. Ales Benda for help with the FCS data analysis. This study was supported by the Czech Science Foundation (17-20613Y to MH). The Group of Adaptive Immunity is supported by an EMBO Installation Grant (3259 to OS) and the Institute of Molecular Genetics of the Czech Academy of Sciences core funding (RVO 68378050). OS is supported by the Purkyne Fellowship provided by the Czech Academy of Sciences. AP and VN are students partially supported by the Faculty of Science, Charles University, Prague.

We acknowledge the Light Microscopy Core Facility, IMG CAS, Prague, Czech Republic, supported by MEYS (LM2015062, CZ.02.1.01/0.0/0.0/16_013/0001775), OPPK (CZ.2.16/3.1.00/21547) and MEYS (LO1419), for their support with obtaining the widefield, and FRAP data presented herein. In addition, we acknowledge Imaging Methods Core Facility at BIOCEV, Prague, Czech Republic, supported by the MEYS CR (Large RI Project LM2018129 Czech-BioImaging) and ERDF (project No. CZ.02.1.01/0.0/0.0/16_013/0001775) for their support with obtaining and analyzing FCS data presented in this paper.

## Author contribution

MH and OS conceived the study and were in charge of the overall direction and planning. AP and MH performed most of the basic and quantitative microscopy and biochemical experiments, MSC performed the expansion microscopy experiments, MSC, KR, MA, and VN performed cloning and cell line generation. PD helped to establish the CRISPR/Cas9 approach. AP, MH and OS wrote the manuscript with the contribution of all the other authors.

## References

1. Forsythe, E. and P.L. Beales, Bardet-Biedl syndrome. Eur J Hum Genet, 2013. 21(1): p. 8–13.

2. Niederlova, V., et al., Meta-analysis of genotype-phenotype associations in Bardet-Biedl syndrome uncovers differences among causative genes. Hum Mutat, 2019. 40(11): p. 2068–2087.

3. Nachury, M.V., et al., A core complex of BBS proteins cooperates with the GTPase Rab8 to promote ciliary membrane biogenesis. Cell, 2007. 129(6): p. 1201–13.

4. Loktev, A.V., et al., A BBSome subunit links ciliogenesis, microtubule stability, and acetylation. Dev Cell, 2008. 15(6): p. 854–65.

5. Wingfield, J.L., K.F. Lechtreck, and E. Lorentzen, Trafficking of ciliary membrane proteins by the intraflagellar transport/BBSome machinery. Essays Biochem, 2018. 62(6): p. 753–763.

6. Ye, F., A.R. Nager, and M.V. Nachury, BBSome trains remove activated GPCRs from cilia by enabling passage through the transition zone. Journal of Cell Biology, 2018. 217(5): p. 1847–1868.

7. Nachury, M.V., The molecular machines that traffic signaling receptors into and out of cilia. Current Opinion in Cell Biology, 2018. 51: p. 124–131.

8. Liu, P.W. and K.F. Lechtreck, The Bardet-Biedl syndrome protein complex is an adapter expanding the cargo range of intraflagellar transport trains for ciliary export. Proceedings of the National Academy of Sciences of the United States of America, 2018. 115(5): p. E934–E943.

9. Jin, H., et al., The conserved Bardet-Biedl syndrome proteins assemble a coat that traffics membrane proteins to cilia. Cell, 2010. 141(7): p. 1208–19.

10. Zhang, Q., et al., BBS proteins interact genetically with the IFT pathway to influence SHH-related phenotypes. Hum Mol Genet, 2012. 21(9): p. 1945–53.

11. Guo, D.F., et al., The BBSome Controls Energy Homeostasis by Mediating the Transport of the Leptin Receptor to the Plasma Membrane. PLoS genetics, 2016. 12(2): p. e1005890.

12. Nishimura, D.Y., et al., Bbs2-null mice have neurosensory deficits, a defect in social dominance, and retinopathy associated with mislocalization of rhodopsin. Proc Natl Acad Sci U S A, 2004. 101(47): p. 16588–93.

13. McIntyre, J.C., M.M. Hege, and N.F. Berbari, Trafficking of ciliary G protein-coupled receptors. Methods Cell Biol, 2016. 132: p. 35–54.

14. Loktev, A.V. and P.K. Jackson, Neuropeptide Y family receptors traffic via the Bardet-Biedl syndrome pathway to signal in neuronal primary cilia. Cell Rep, 2013. 5(5): p. 1316–29.

15. Berbari, N.F., et al., Bardet-Biedl syndrome proteins are required for the localization of G protein-coupled receptors to primary cilia. Proceedings of the National Academy of Sciences of the United States of America, 2008. 105(11): p. 4242–6.

16. Woodsmith, J., et al., Protein interaction perturbation profiling at amino-acid resolution. Nat Methods, 2017. 14(12): p. 1213–1221.

17. Katoh, Y., et al., Architectures of multisubunit complexes revealed by a visible immunoprecipitation assay using fluorescent fusion proteins. J Cell Sci, 2015. 128(12): p. 2351-62.

18. Klink, B.U., et al., A recombinant BBSome core complex and how it interacts with ciliary cargo. Elife, 2017. 6.

19. Mourao, A., et al., Structural basis for membrane targeting of the BBSome by ARL6. Nat Struct Mol Biol, 2014. 21(12): p. 1035–41.

20. Knockenhauer, K.E. and T.U. Schwartz, Structural Characterization of Bardet-Biedl Syndrome 9 Protein (BBS9). J Biol Chem, 2015. 290(32): p. 19569–83.

21. Zhang, Q., et al., Intrinsic protein-protein interaction-mediated and chaperonin-assisted sequential assembly of stable bardet-biedl syndrome protein complex, the BBSome. The Journal of biological chemistry, 2012. 287(24): p. 20625–35.

22. Klink, B.U., et al., Structure of the human BBSome core complex. Elife, 2020. 9.

23. Chou, H.T., et al., The Molecular Architecture of Native BBSome Obtained by an Integrated Structural Approach. Structure, 2019. 27(9): p. 1384–1394 e4.

24. Singh, S.K., et al., Structure and activation mechanism of the BBSome membrane protein trafficking complex. Elife, 2020. 9.

25. Huranova, M., et al., The differential interaction of snRNPs with pre-mRNA reveals splicing kinetics in living cells. The Journal of cell biology, 2010. 191(1): p. 75–86.

26. Huranova, M., et al., Dynamic assembly of the exomer secretory vesicle cargo adaptor subunits. EMBO reports, 2016. 17(2): p. 202–19.

27. Labun, K., et al., CHOPCHOP v3: expanding the CRISPR web toolbox beyond genome editing. Nucleic Acids Res, 2019. 47(W1): p. W171–W174.

28. Ran, F.A., et al., Genome engineering using the CRISPR-Cas9 system. Nat Protoc, 2013. 8(11): p. 2281–2308.

29. Gambarotto, D., et al., Imaging cellular ultrastructures using expansion microscopy (U-ExM). Nat Methods, 2019. 16(1): p. 71–74.

30. Schwille, P., et al., Molecular dynamics in living cells observed by fluorescence correlation spectroscopy with one- and two-photon excitation. Biophys J, 1999. 77(4): p. 2251–65.

31. Widengren, J., U. Mets, and R. Rigler, Fluorescence correlation spectroscopy of triplet states in solution: a theoretical and experimental study. The Journal of Physical Chemistry, 1995. 99(36): p. 13368–13379.

32. Kremers, G.J., et al., Cyan and yellow super fluorescent proteins with improved brightness, protein folding, and FRET Forster radius. Biochemistry, 2006. 45(21): p. 6570–80.

33. Hernandez-Hernandez, V., et al., Bardet-Biedl syndrome proteins control the cilia length through regulation of actin polymerization. Human molecular genetics, 2013. 22(19): p. 3858–68.

34. Kim, J.C., et al., The Bardet-Biedl protein BBS4 targets cargo to the pericentriolar region and is required for microtubule anchoring and cell cycle progression. Nature Genetics, 2004. 36(5): p. 462–470.

35. Zhang, Q., et al., BBS7 is required for BBSome formation and its absence in mice results in Bardet-Biedl syndrome phenotypes and selective abnormalities in membrane protein trafficking. Journal of cell science, 2013. 126(Pt 11): p. 2372–80.

36. Quarantotti, V., et al., Centriolar satellites are acentriolar assemblies of centrosomal proteins. EMBO J, 2019. 38(14): p. e101082.

37. Dammermann, A. and A. Merdes, Assembly of centrosomal proteins and microtubule organization depends on PCM-1. J Cell Biol, 2002. 159(2): p. 255–66.

38. Chamling, X., et al., The centriolar satellite protein AZI1 interacts with BBS4 and regulates ciliary trafficking of the BBSome. PLoS Genet, 2014. 10(2): p. e1004083.

39. Kim, J.C., et al., The Bardet-Biedl protein BBS4 targets cargo to the pericentriolar region and is required for microtubule anchoring and cell cycle progression. Nature genetics, 2004. 36(5): p. 462–70.

40. Stowe, T.R., et al., The centriolar satellite proteins Cep72 and Cep290 interact and are required for recruitment of BBS proteins to the cilium. Mol Biol Cell, 2012. 23(17): p. 3322–35.

41. Seo, S., et al., A novel protein LZTFL1 regulates ciliary trafficking of the BBSome and Smoothened. PLoS Genet, 2011. 7(11): p. e1002358.

42. Marion, V., et al., Exome sequencing identifies mutations in LZTFL1, a BBSome and smoothened trafficking regulator, in a family with Bardet--Biedl syndrome with situs inversus and insertional polydactyly. J Med Genet, 2012. 49(5): p. 317–21.

43. Jiang, J., et al., Depletion of BBS Protein LZTFL1 Affects Growth and Causes Retinal Degeneration in Mice. J Genet Genomics, 2016. 43(6): p. 381–91.

44. Seo, S., et al., BBS6, BBS10, and BBS12 form a complex with CCT/TRiC family chaperonins and mediate BBSome assembly. Proceedings of the National Academy of Sciences of the United States of America, 2010. 107(4): p. 1488–93.

45. Stoetzel, C., et al., Identification of a novel BBS gene (BBS12) highlights the major role of a vertebrate-specific branch of chaperonin-related proteins in Bardet-Biedl syndrome. Am J Hum Genet, 2007. 80(1): p. 1–11.

46. Stoetzel, C., et al., BBS10 encodes a vertebrate-specific chaperonin-like protein and is a major BBS locus. Nat Genet, 2006. 38(5): p. 521–4.

47. Katsanis, N., et al., Mutations in MKKS cause obesity, retinal dystrophy and renal malformations associated with Bardet-Biedl syndrome. Nat Genet, 2000. 26(1): p. 67–70.

